# A Root Foundation Model for Zero-Shot Segmentation

**DOI:** 10.64898/2026.05.14.725129

**Authors:** Abraham George Smith, Sotiris Lamprinidis, Agata Wlaszczyk, Jens Petersen

## Abstract

Foundation models pre-trained on massive datasets have demonstrated impressive performance, but in some specialised domains have been found to have lower accuracy. Domain-specific foundation models target a particular domain such as retinal or plant images. These domain-specific models have shown inconsistent results and the benefit to root segmentation is unknown. We train and evaluate the first domain-specific foundation model for root segmentation. Evaluation uses a leave-one-dataset-out design across nine diverse root datasets with two architectures. Applied zero-shot to unseen datasets, the root foundation model achieves 92% of fine-tuned Dice on average (0.636 versus 0.698), with 5 of 9 datasets above 90%. With 10 patches of few-shot fine-tuning, the root foundation model recovers 95% of its full-data Dice on average, versus 69% for a general pre-trained model. At low patch counts the general pre-trained model often failed to converge, with 5 of 9 datasets giving Dice below 0.05 at 3 patches, while the root foundation model produced Dice above 0.47 on every dataset and patch count. With full target-data fine-tuning, the two perform comparably, with mean improvements of +0.011 Dice for MobileSAM and +0.022 for M2F Swin-S, neither significant (Wilcoxon *p* = 0.150 and 0.064). We release our pre-trained MobileSAM root foundation model for use with RootPainter, enabling fully automatic root segmentation on new datasets with an ordinary laptop or desktop computer, with no need for annotation or training.

## Introduction

Deep learning has transformed root image analysis, with convolutional and Transformer-based architectures achieving high segmentation accuracy across diverse imaging conditions [Smith et al., 2020, Baykalov et al., 2023, Delory et al., 2022]. Interactive tools such as RootPainter [Smith et al., 2022] have enabled researchers to apply deep learning segmentation to root phenotyping across various species and conditions [Sell et al., 2022, Chen et al., 2022, Malinowska et al., 2022]. However, each new imaging setup often requires training a new model on a manually annotated subset of the experimental data [e.g., Bauer et al., 2022, Callwood et al., 2025, Fagnant et al., 2025]. Banet et al. [2024] showed that models trained at one location were not suitable for direct application at a separate location, with cross-site RMSE increasing by 20–74%. Manual annotation of root images is also time-consuming. Per-pixel labelling of chicory roots in rhizobox images of 300 × 170 mm takes around 30 minutes per image [Smith et al., 2020], while root tracing in Rootfly [Zeng et al., 2008] of 15 × 15 mm forest minirhizotron windows took 80 hours for 2,116 images [Handy et al., 2024]. Recent work comparing 21 architectures across nine root datasets found that pre-trained models significantly outperform models trained from scratch, with Transformers benefiting more from pre-training than ConvNets [Smith et al., 2026]. MobileSAM, a lightweight adaptation of the Segment Anything Model [Kirillov et al., 2023, Zhang et al., 2023], achieved the highest segmentation accuracy while maintaining computational efficiency.

These findings raise a natural question: if general pre-training (on ImageNet, SAM, or similar large-scale datasets) improves root segmentation, would domain-specific pre-training on root images provide further benefit?

### Contributions

1. The first root-specific foundation model for segmentation
2. Zero-shot segmentation of unseen root datasets at 92% of fine-tuned Dice (above 90% for 5 of 9 datasets)
3. A leave-one-dataset-out evaluation across nine datasets showing that root-specific pre-training substantially exceeds general pre-training under low-data fine-tuning
4. Release of the trained MobileSAM root foundation model for use with RootPainter on CPU, enabling fully automatic root segmentation on an ordinary laptop or desktop computer

To investigate this, we use a leave-one-dataset-out crossvalidation design across nine root image datasets. For each fold, we train a foundation model on eight datasets and evaluate on the ninth, comparing foundation model performance (both zero-shot and fine-tuned) against baselines that skip foundation training entirely.

## Related Work

### Continued pre-training

In natural language processing, Gururangan et al. [2020] studied continued pre-training, where a general pretrained model is trained further on domain-specific data. They found continued pre-training improves target task performance, with larger improvements when the domain differs more from the original pre-training data.

### Domain-specific foundation models in computer vision

In computer vision, Isztl et al. [2025] compared a retinal foundation model against ImageNet-pre-trained models. Domain-specific pre-training only helped for the most challenging task (diabetic retinopathy grading). For easier tasks, general-purpose models performed equally well or better. Similarly, Hou et al. [2025] found that a general-purpose model outperformed a retinal-specific model for ocular disease detection, but the domain-specific model performed better for systemic disease prediction. Zhou et al. [2025] found that while domain-specific pretraining still outperformed generalist models when finetuned, the performance difference was small (AUROC 0.830 vs 0.816). A consistent finding across these studies is that pre-trained initialisation, whether general or domain-specific, significantly outperforms training from scratch [Isztl et al., 2025].

### Zero-shot segmentation with foundation models

Zero-shot performance of general foundation models has been found to be poor in several specialised domains. The Dice of zero-shot SAM was only 24–29% of fine-tuned performance on retinal vessel segmentation [Shi et al., 2023], and low zero-shot Dice scores have been reported for scientific microscopy (0.17) [Mukherjee et al., 2025] and remote sensing (mIoU 0.10–0.24) [Zhang et al., 2024]. In contrast, Chattopadhyay et al. [2025] found that foundation models trained on domain-specific data achieved zeroshot performance close to fine-tuned accuracy on held-out datasets within the same domain.

### Foundation models for plant phenotyping

For plant phenotyping, Chen et al. [2023] showed that general vision foundation models can be efficiently adapted to tasks like leaf counting and segmentation, achieving performance close to task-specific models. For root image analysis, Smith et al. [2026] found that pre-trained models significantly outperformed models trained from scratch. However, both studies used only general pre-training. To the best of our knowledge, no foundation model trained on root images has been developed. In this work, we create the first root foundation model and evaluate whether it provides benefit over general pre-training alone.

### Hypotheses

Given that domain-specific pre-training shows inconsistent benefits across vision domains, and that no root-specific foundation model has been tested, we hypothesise:

- Fine-tuning a root-specific foundation model trained on diverse root datasets achieves higher accuracy than fine-tuning a general foundation model, demonstrated by higher Dice across the nine held-out datasets.
- Fine-tuning the root foundation model achieves higher accuracy than applying it zero-shot, demonstrated by higher Dice across the nine held-out datasets.

## Methods

Throughout this paper, we use the following terminology. *General pre-training* refers to the original model weights trained on large-scale non-root datasets (e.g. ImageNet, SA-1B). *Root foundation training* refers to the intermediate stage of supervised segmentation training on eight root datasets. *Fine-tuning* refers to supervised segmentation training on the target (held-out) dataset.

Where applicable, we follow the experimental setup of Smith et al. [2026] (datasets, architectures, training protocol, learning rate) to enable direct comparison with their baselines.

### Datasets

We use nine publicly available root image datasets, spanning diverse species and imaging modalities (Figure 1). Annotation approaches also differ: DeepRootLab uses corrective annotations, while the remaining datasets use dense pixel-level annotations of all visible roots.

**Fig. 1:**
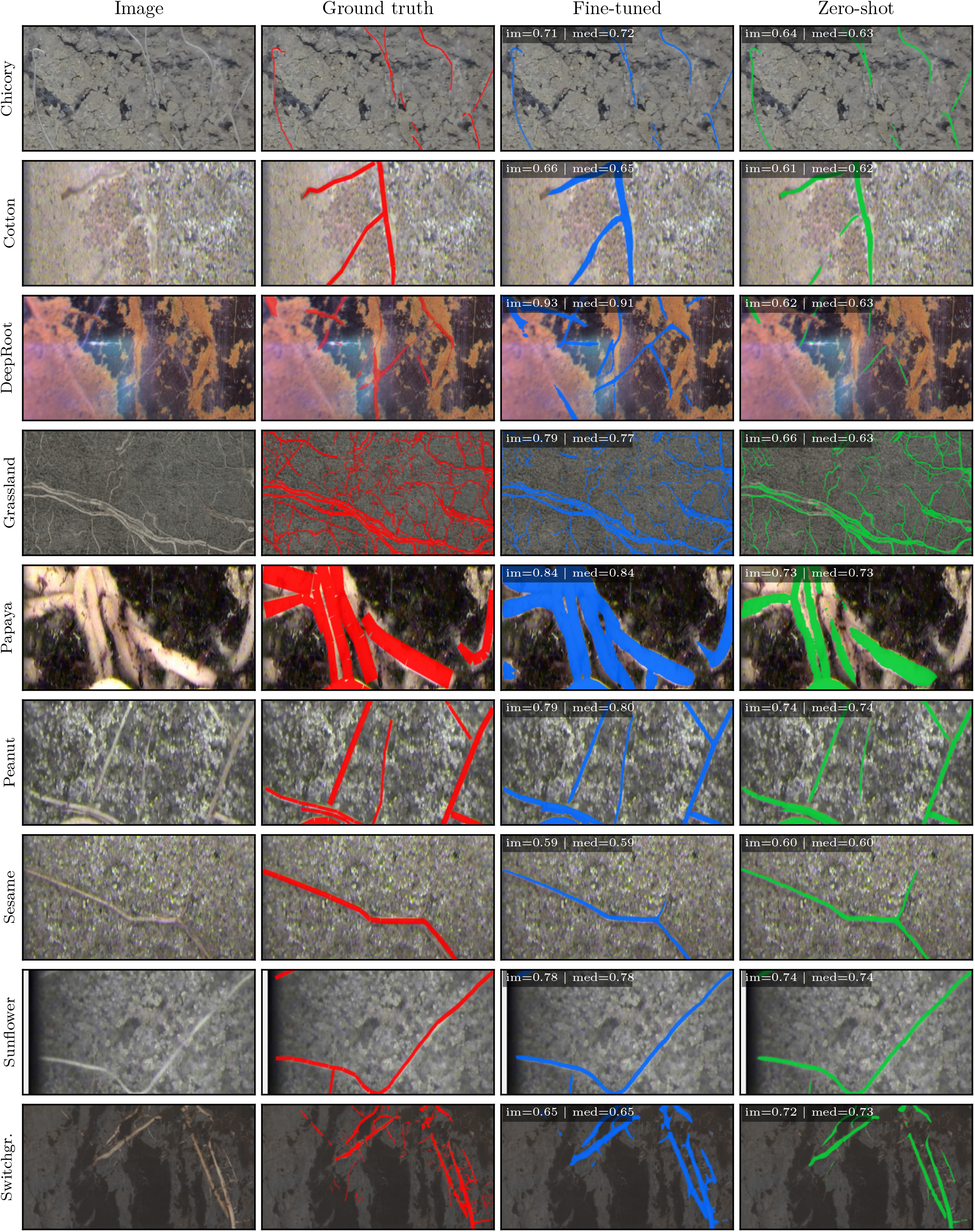
For MobileSAM (general pre-trained weights, leave-one-out, test split), columns show the original image, the ground truth overlay (red), the fine-tuned prediction (blue), and the zero-shot prediction (green). For each dataset, the shown image is selected from those with above-average fine-tuned prediction pixel count, choosing the one that minimises the joint deviation of its fine-tuned and zero-shot Dice from the dataset’s medians, |FT −FT_med_ |+ |ZS−ZS_med_| . Text overlays on the fine-tuned and zero-shot tiles show this image’s Dice (im) and the dataset’s median Dice (med). All rows are centre-cropped to a 2.5:1 aspect ratio for row-height consistency, Chicory is additionally zoomed 4× to the most root-dense region of the ground truth, and Switchgrass is rotated 90 degrees. DeepRoot uses corrective annotations (red = foreground, green = background).

### DeepRootLab

Images of eleven species collected at the DeepRootLab facility [Han et al., 2026], which includes 48 plots with 144 6-metre-long minirhizotron tubes.

#### Grassland

Collected in alpine meadows [Möhl et al., 2022], capturing natural root-soil interactions under field conditions.

#### Chicory

Densely annotated rhizobox images from a controlled setup [Smith et al., 2020, 2019, Thorup-Kristensen et al., 2020].

#### PRMI Collection

Six minirhizotron datasets, namely Papaya, Peanut, Sesame, Sunflower, Cotton, and Switchgrass, released as part of the PRMI benchmark [Xu et al., 2022].

Dataset sizes vary substantially, from 48 images (Chicory) to 19,625 (Peanut), motivating the balanced sampling strategy described below.

### Model architectures

We run all experiments with MobileSAM and M2F Swin-S, which achieved the highest and second-highest Dice scores respectively in a previous comparison of 21 architectures evaluated for root segmentation [Smith et al., 2026]:

**MobileSAM** [Zhang et al., 2023] uses a Vision Transformer encoder (ViT-Tiny, 10.1M parameters) distilled from the original SAM model [Kirillov et al., 2023].

**Mask2Former with Swin-S** (M2F Swin-S, 68.7M parameters) pairs a Swin Transformer encoder [Liu et al., 2021] with a Mask2Former decoder [Cheng et al., 2022].

### Experimental design

We use a leave-one-dataset-out cross-validation design with nine folds. For each fold, one dataset serves as the held-out test set, and the remaining eight datasets are used for foundation model training.

For each fold, we compare three conditions (Table I):

**TABLE I:**
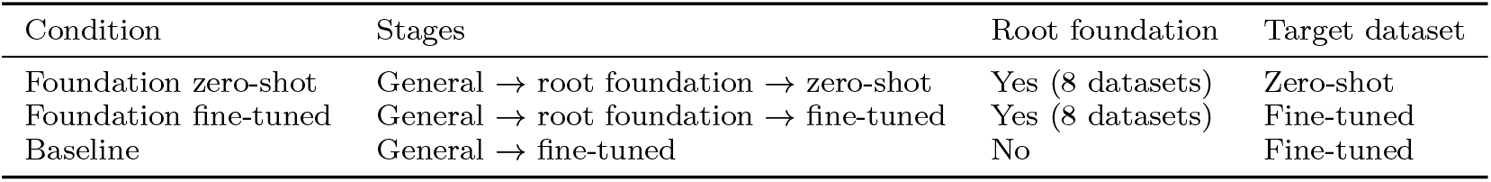
Experimental conditions for each cross-validation fold.

| Condition | Stages | Root foundation | Target dataset |
| --- | --- | --- | --- |
| Foundation zero-shot | General → root foundation → zero-shot | Yes (8 datasets) | Zero-shot |
| Foundation fine-tuned | General → root foundation → fine-tuned | Yes (8 datasets) | Fine-tuned |
| Baseline | General → fine-tuned | No | Fine-tuned |

The Baseline condition skips root-specific foundation training entirely, fine-tuning directly on the target dataset from general pre-trained weights.

#### Combined training

In addition to the leave-one-out design, we train a model on all nine datasets simultaneously using the same balanced sampling and training protocol. This model is evaluated on each dataset’s validation and test splits without further fine-tuning. Because the model has seen training data from all nine datasets, this is the model we release for others to use, trained on all available root datasets.

### Foundation model training

For each cross-validation fold, we train a foundation model on the combined training sets of eight datasets.

#### Balanced sampling

Dataset sizes range from 48 images for Chicory to 19,625 for Peanut, a factor of more than 400 between them. To prevent large datasets from dominating training, we use balanced sampling, where each training epoch draws an equal number of samples from each dataset. Specifically, the epoch size equals 8*×* the size of the smallest dataset, with the same number of samples drawn from each dataset for each epoch. This ensures the model sees diverse training data rather than overfitting to the dataset with the largest number of images.

#### Training protocol

Training uses the AdamW optimiser with a learning rate of 0.0001, 16-bit mixed precision, and gradient norm clipping at a maximum norm of 1. A warm-up schedule of 2 epochs is applied, followed by early stopping if validation Dice does not improve for 20 epochs. The loss function is an equal-weighted sum of Dice loss and binary cross-entropy. Batches containing no root annotations are excluded. During root foundation training, we hold out a validation set from each of the eight training datasets and monitor mean validation Dice across all eight.

### Target dataset evaluation

#### Zero-shot

The foundation model is applied directly to the held-out test set without any training on the target dataset.

#### Fine-tuned

The foundation model is fine-tuned on the held-out dataset’s training set, then evaluated on its test set. Fine-tuning uses the same protocol as foundation training (AdamW, early stopping).

#### Baselines

The Baseline starts from general pre-trained weights (see Model architectures) and skips root-specific foundation training.

### Few-shot fine-tuning

To assess how the root foundation model behaves under low-data fine-tuning, we fine-tuned MobileSAM with 1, 2, 3, 5, 10, 20, or 50 patches of 1024 × 1024 pixels sampled from each held-out dataset’s training images. We compared two starting points, the leave-one-out root foundation model and general pre-trained weights, with all other training settings matching the full fine-tuning protocol. The full-dataset training set is also expressed in patches, summing *W* /1024 *H*/1024 across all training images. Validation Dice was used for early stopping during fine-tuning, and the reported scores are Dice on the heldout test set.

### Evaluation metrics

#### Dice coefficient

Primary segmentation quality metric. True positives, false positives, and false negatives are summed across all images in a dataset before computing the Dice score, rather than averaging per-image scores.

#### Root length correlation

Secondary metric capturing agreement between predicted and ground-truth root length per image. Total root length is extracted from each segmentation using RhizoVision Explorer [Seethepalli et al., 2021], and Pearson correlation *r* between predicted and ground-truth lengths is computed across all images in a dataset. DeepRoot is excluded from this metric because its annotations are corrective rather than dense, often leaving roots only partially annotated.

### Statistical analysis

We compare conditions using Wilcoxon signed-rank tests across the cross-validation folds, with each fold providing one paired observation per condition. Wilcoxon signed-rank is appropriate here because the sample size is small (*n* ≤ 9) and Dice coefficients are bounded between 0 and 1, making normality assumptions for parametric tests questionable. We use one-sided alternatives, testing whether root foundation training improves over the baseline and whether fine-tuning improves over zero-shot.

### Computational resources

All training is performed on machines equipped with an AMD Ryzen 9 7950X CPU, 32GB RAM, and an NVIDIA RTX 4090 GPU with 24GB VRAM.

## Results

Results reported in this section are on the **test** split.

### Does foundation training help?

For MobileSAM (Table III), root foundation training gives a mean improvement of +0.011 Dice, with 7 of 9 datasets showing positive deltas. The Wilcoxon test does not reach significance (Wilcoxon *W* = 32, *n* = 9, *p* =0.150).

For M2F Swin-S (Table III), the mean delta is +0.022 Dice, with DeepRoot the largest at +0.139. 7 of 9 datasets show positive deltas, with Papaya and Sunflower negative. The test is not significant (*W* = 36, *n* = 9, *p* = 0.064). Root length correlation shows no benefit from foundation training when fine-tuning on the full labelled dataset (Table IV), with means of -0.002 (MobileSAM) and +0.000 (M2F Swin-S).

### How much does fine-tuning add?

Table V compares the root foundation model fine-tuned to the target dataset against the same model applied zeroshot (i.e. without target-dataset fine-tuning). Sixteen of eighteen per-dataset deltas are positive (nine datasets × two architectures), ranging up to +0.193 for MobileSAM on DeepRoot and +0.178 for M2F on Chicory. The two negative deltas are MobileSAM on Switchgrass at − 0.058 and M2F on Sesame at − 0.006. The Wilcoxon signedrank test gave *W* = 40, *p* = 0.020 for MobileSAM and *W* = 43, *p* = 0.006 for M2F Swin-S, each with *n* = 9. Length r deltas (Table VI) are smaller. Length r is higher for fine-tuned than zero-shot in thirteen of sixteen comparisons (eight datasets and two architectures, excluding DeepRoot). The exceptions are MobileSAM on Sesame and Sunflower, and M2F on Sesame.

### Zero-shot performance

The MobileSAM root foundation model applied zeroshot achieves 72–108% of its fine-tuned Dice depending on the dataset (108% means zero-shot exceeded fine-tuned on Switchgrass), with a mean of 92% (Figure 2). For Sesame, Sunflower, and Cotton, the gap is small at 0, 2, and 3% relative difference respectively, and for Switchgrass, zeroshot exceeds fine-tuned by 8%. For DeepRoot, the drop is more substantial at 28% relative difference. Figure 1 shows the visual similarity between the zero-shot and fine-tuned model segmentations.

**Fig. 2:**
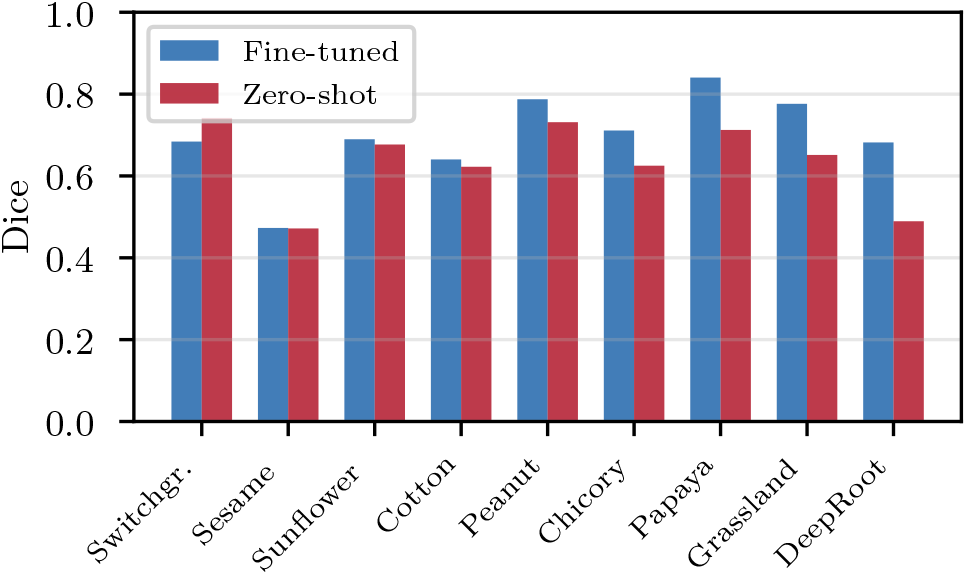
Fine-tuned vs zero-shot Dice for MobileSAM with general pre-trained weights (leave-one-out, test split). Datasets are sorted by the size of the gap.

Root length correlation with ground truth is less affected than Dice by the switch from fine-tuned to zero-shot (Table VII). Zero-shot is on average 99% of fine-tuned root length *r*, compared to 92% for Dice. For Sunflower and Sesame, zero-shot root length correlation slightly exceeds fine-tuned (Sunflower: 0.935 vs 0.886; Sesame: 0.947 vs 0.912), and for Chicory, Peanut, and Grassland, zero-shot is 98–99% of fine-tuned.

### Few-shot fine-tuning

Figure 3 shows test Dice as a function of the number of training patches used for fine-tuning, comparing the root foundation model and a general pre-trained model as starting points. Each method is compared against its own full-data Dice. Fine-tuning the root foundation model on 10 patches recovers 71–103% of its full-data Dice (mean 95%), while fine-tuning a general pre-trained model on the same 10 patches recovers only 1–95% of its full-data Dice (mean 69%). Increasing to 50 patches narrows the gap, with the general pre-trained model reaching 59–102% (mean 90%) while the root foundation model reaches 73–101% (mean 96%). Ten patches represents between 0.09% (Peanut) and 4.31% (Chicory) of each dataset’s full training set, with a mean of 1.46%.

**Fig. 3:**
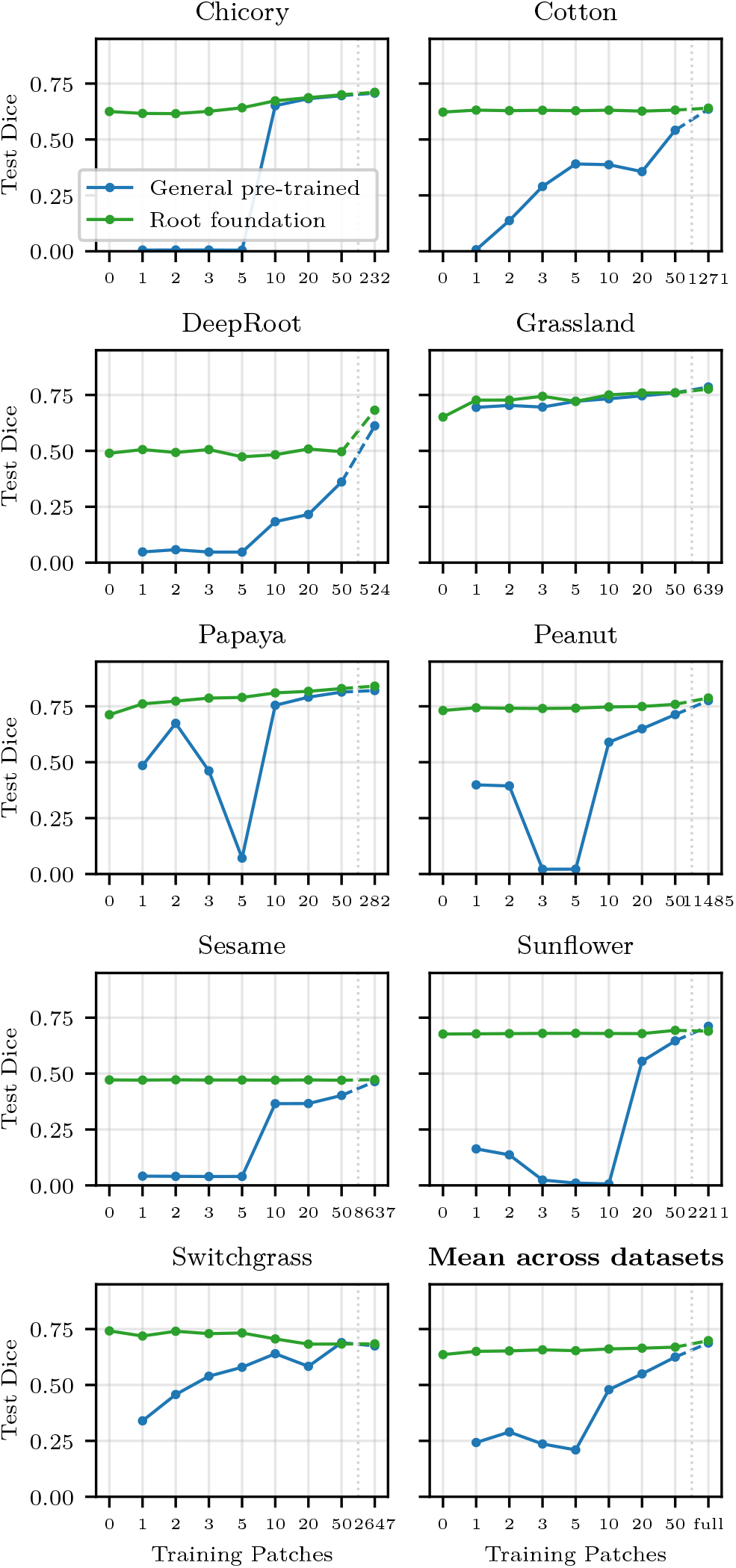
Test Dice for MobileSAM fine-tuned on small numbers of training patches (1, 2, 3, 5, 10, 20, 50), starting from either the root foundation model (green) or general pre-trained weights (blue). The 0 column shows zeroshot Dice from the root foundation model. The rightmost column shows fine-tuning on the full training set, with the per-dataset patch count given on the x-axis. The dashed segment indicates that the gap between 50 patches and the full training set is not to scale.

Fine-tuning from the general pre-trained model frequently failed to converge at low patch counts. At 3 patches, five of nine datasets produced Dice below 0.05, and at 5 patches, six of nine produced Dice below 0.10 (Figure 3). Fine-tuning from the root foundation model produced Dice above 0.47 on every dataset at every patch count we tested, suggesting that reducing the domain gap improves robustness to limited fine-tuning data.

### Combined training

In addition to the leave-one-out experiments, we train a combined model on all nine datasets. Training on all nine datasets at once gives a mean Dice of 0.700 for MobileSAM and 0.694 for M2F, comparable to per-dataset fine-tuning at 0.698 and 0.696 respectively. We release this combined model rather than a dataset-specific model, as we expect it to generalise better to unseen root datasets.

## Discussion

Our first hypothesis was that root foundation training would improve over general pre-training when fine-tuned on a target dataset. The mean delta was +0.011 Dice for MobileSAM and +0.022 for M2F Swin-S, neither reaching significance (*p* = 0.150 and 0.064; Table III). Our second hypothesis was that fine-tuning would improve over zeroshot. This was supported for both architectures (MobileSAM *p* = 0.020; M2F Swin-S *p* = 0.006; Table V), with zero-shot performance averaging 92% of fine-tuned Dice.

### When domain-specific foundation models help

Root foundation training before fine-tuning shows inconsistent benefits across the nine datasets and two architectures. DeepRootLab has the highest mean delta across the two architectures at +0.104, with values of +0.070 for MobileSAM and +0.139 for M2F Swin-S in Table III. DeepRootLab is also the most heterogeneous of the nine datasets, imaged in agricultural field conditions across multiple crop species and seasons, while the other datasets use more controlled minirhizotron or rhizobox setups with a single species each. Root foundation training also provides a substantial benefit when fine-tuning data is limited. Figure 3 shows that with only 10 patches the root foundation model recovers 95% of full-data Dice on average versus 69% for general pre-training. In retinal imaging, Isztl et al. [2025] found that a domain-specific model outperformed general models only on diabetic retinopathy grading, the most challenging task in their evaluation. Hou et al. [2025] reported that a generalpurpose model outperformed the domain-specific model on ocular disease detection, which they describe as identifying well-defined lesions with distinct visual patterns, but the domain-specific model performed better on systemic disease prediction such as heart failure and stroke. Across these studies, including ours, domain-specific foundation models appear to provide greater benefit on challenging and complex datasets or when labelled data is limited.

### Architecture comparison under root foundation training

In our test-split evaluation, MobileSAM with root foundation training reached a mean Dice of 0.698, slightly above its 0.687 baseline. M2F Swin-S reached 0.696 with root foundation training and 0.674 without (Table II). Neither lift was statistically significant. The two architectures end up at essentially the same Dice once root foundation training is added, with MobileSAM remaining the simpler model at 10.1M parameters versus 68.7M for M2F Swin-S [Smith et al., 2026].

**TABLE II:** Mean Dice *±* SD and root length correlation (test split) across cross-validation folds. Root length *r* = Pearson correlation between predicted and ground-truth total root length (8 datasets; DeepRoot excluded because its annotations are partial).

| Condition | Dice | | Root length $r$ | |
| --- | --- | --- | --- | --- |
|  | MobileSAM | M2F Swin-S | MobileSAM | M2F Swin-S |
| Foundation zero-shot (root foundation, no fine-tuning) | 0.636 ± 0.098 | 0.620 ± 0.097 | 0.930 ± 0.035 | 0.922 ± 0.038 |
| Foundation fine-tuned (root foundation, with fine-tuning) | **0.698 ± 0.106** | **0.696 ± 0.110** | 0.940 ± 0.036 | 0.946 ± 0.025 |
| Baseline (general pre-trained, with fine-tuning) | 0.687 ± 0.109 | 0.674 ± 0.126 | **0.942 ± 0.036** | **0.946 ± 0.026** |

**TABLE III:** Per-dataset Dice with vs without root foundation training, both from general pre-trained weights. Positive Δ means foundation training helped.

| Model | Dataset | Baseline | Foundation | $\Delta$ |
| --- | --- | --- | --- | --- |
| MobileSAM | Chicory | 0.707 | 0.711 | +0.004 |
|  | Cotton | 0.636 | 0.640 | +0.005 |
|  | DeepRoot | 0.612 | 0.682 | +0.070 |
|  | Grassland | 0.786 | 0.776 | -0.009 |
|  | Papaya | 0.821 | 0.840 | +0.020 |
|  | Peanut | 0.776 | 0.787 | +0.012 |
|  | Sesame | 0.464 | 0.473 | +0.009 |
|  | Sunflower | 0.711 | 0.690 | -0.022 |
|  | Switchgr. | 0.675 | 0.684 | +0.009 |
|  | Mean | 0.687 | 0.698 | +0.011 |
| M2F Swin-S | Chicory | 0.712 | 0.719 | +0.007 |
|  | Cotton | 0.589 | 0.620 | +0.032 |
|  | DeepRoot | 0.523 | 0.662 | +0.139 |
|  | Grassland | 0.773 | 0.791 | +0.018 |
|  | Papaya | 0.840 | 0.829 | -0.011 |
|  | Peanut | 0.771 | 0.773 | +0.002 |
|  | Sesame | 0.459 | 0.463 | +0.004 |
|  | Sunflower | 0.683 | 0.670 | -0.013 |
|  | Switchgr. | 0.713 | 0.732 | +0.019 |
|  | Mean | 0.674 | 0.696 | +0.022 |

**TABLE IV:** Per-dataset root length correlation (*r*) with vs without root foundation training. DeepRoot is excluded because its annotations are partial.

| Model | Dataset | Baseline | Foundation | $\Delta$ |
| --- | --- | --- | --- | --- |
| MobileSAM | Chicory | 0.998 | 0.998 | 0.000 |
|  | Cotton | 0.953 | 0.959 | +0.006 |
|  | Grassland | 0.960 | 0.949 | -0.011 |
|  | Papaya | 0.894 | 0.923 | +0.028 |
|  | Peanut | 0.971 | 0.968 | -0.003 |
|  | Sesame | 0.901 | 0.912 | +0.011 |
|  | Sunflower | 0.941 | 0.886 | -0.055 |
|  | Switchgr. | 0.916 | 0.924 | +0.008 |
|  | Mean | 0.942 | 0.940 | -0.002 |
| M2F Swin-S | Chicory | 0.998 | 0.999 | +0.001 |
|  | Cotton | 0.960 | 0.935 | -0.025 |
|  | Grassland | 0.936 | 0.944 | +0.008 |
|  | Papaya | 0.926 | 0.927 | +0.001 |
|  | Peanut | 0.962 | 0.967 | +0.006 |
|  | Sesame | 0.929 | 0.934 | +0.005 |
|  | Sunflower | 0.944 | 0.941 | -0.004 |
|  | Switchgr. | 0.916 | 0.923 | +0.007 |
|  | Mean | 0.946 | 0.946 | -0.000 |

**TABLE V:** Fine-tuned vs zero-shot Dice for the root foundation model with general pre-trained weights. Positive Δ means fine-tuning helped.

| Model | Dataset | Zero-shot | Fine-tuned | $\Delta$ |
| --- | --- | --- | --- | --- |
| MobileSAM | Chicory | 0.625 | 0.711 | +0.086 |
|  | Cotton | 0.622 | 0.640 | +0.018 |
|  | DeepRoot | 0.489 | 0.682 | +0.193 |
|  | Grassland | 0.651 | 0.776 | +0.125 |
|  | Papaya | 0.712 | 0.840 | +0.128 |
|  | Peanut | 0.731 | 0.787 | +0.056 |
|  | Sesame | 0.472 | 0.473 | +0.001 |
|  | Sunflower | 0.677 | 0.690 | +0.013 |
|  | Switchgr. | 0.742 | 0.684 | -0.058 |
|  | Mean | 0.636 | 0.698 | +0.062 |
| M2F Swin-S | Chicory | 0.541 | 0.719 | +0.178 |
|  | Cotton | 0.604 | 0.620 | +0.016 |
|  | DeepRoot | 0.506 | 0.662 | +0.156 |
|  | Grassland | 0.628 | 0.791 | +0.162 |
|  | Papaya | 0.726 | 0.829 | +0.103 |
|  | Peanut | 0.726 | 0.773 | +0.048 |
|  | Sesame | 0.469 | 0.463 | -0.006 |
|  | Sunflower | 0.669 | 0.670 | +0.001 |
|  | Switchgr. | 0.711 | 0.732 | +0.021 |
|  | Mean | 0.620 | 0.696 | +0.076 |

**TABLE VI:** Per-dataset root length correlation (*r*), finetuned vs zero-shot. DeepRoot is excluded because its annotations are partial.

| Model | Dataset | Zero-shot | Fine-tuned | $\Delta$ |
| --- | --- | --- | --- | --- |
| MobileSAM | Chicory | 0.980 | 0.998 | +0.018 |
|  | Cotton | 0.927 | 0.959 | +0.032 |
|  | Grassland | 0.941 | 0.949 | +0.008 |
|  | Papaya | 0.861 | 0.923 | +0.061 |
|  | Peanut | 0.946 | 0.968 | +0.022 |
|  | Sesame | 0.947 | 0.912 | -0.035 |
|  | Sunflower | 0.935 | 0.886 | -0.049 |
|  | Switchgr. | 0.901 | 0.924 | +0.023 |
|  | Mean | 0.930 | 0.940 | +0.010 |
| M2F Swin-S | Chicory | 0.953 | 0.999 | +0.046 |
|  | Cotton | 0.916 | 0.935 | +0.019 |
|  | Grassland | 0.914 | 0.944 | +0.030 |
|  | Papaya | 0.914 | 0.927 | +0.013 |
|  | Peanut | 0.953 | 0.967 | +0.015 |
|  | Sesame | 0.946 | 0.934 | -0.012 |
|  | Sunflower | 0.938 | 0.941 | +0.003 |
|  | Switchgr. | 0.839 | 0.923 | +0.084 |
|  | Mean | 0.922 | 0.946 | +0.025 |

**TABLE VII:** Per-dataset root length correlation *r* for the root foundation model fine-tuned vs zero-shot (MobileSAM, leave-one-out, test split). Mean is 99% of finetuned across the 8 datasets shown. Per-dataset Dice values are shown in Figure 2. DeepRoot is excluded because its annotations are partial.

| Dataset | Fine-tuned | Zero-shot | % |
| --- | --- | --- | --- |
| Sunflower | 0.886 | 0.935 | 105% |
| Sesame | 0.912 | 0.947 | 104% |
| Grassland | 0.949 | 0.941 | 99% |
| Chicory | 0.998 | 0.980 | 98% |
| Peanut | 0.968 | 0.946 | 98% |
| Switchgr. | 0.924 | 0.901 | 97% |
| Cotton | 0.959 | 0.927 | 97% |
| Papaya | 0.923 | 0.861 | 93% |
| Mean | 0.940 | 0.930 | 99% |

### Zero-shot segmentation is close to fine-tuned accuracy

Fine-tuning on the target data improves over zeroshot on most datasets for both architectures (Table V), with two exceptions, MobileSAM on Switchgrass where zero-shot exceeds fine-tuned by 0.058 Dice, and M2F on Sesame where zero-shot exceeds fine-tuned by 0.006 Dice. In many cases zero-shot performance is close enough to be useful, with over 90% of fine-tuned Dice for 5 of 9 datasets and 92% on average. When a general-purpose model is applied zero-shot to a specialised domain, performance can be poor. Shi et al. [2023] found that zeroshot SAM was only 24–29% of fine-tuned Dice on retinal vessel segmentation, a task that, like root segmentation, requires delineating continuous branching structures. Most evaluations of domain-specific foundation models focus on whether they improve fine-tuned performance over general pre-training [Isztl et al., 2025, Hou et al., 2025]. However, Chattopadhyay et al. [2025] found that domain-specific foundation models achieved zero-shot performance close to fine-tuned accuracy on held-out datasets within the same domain. Our results support this finding for root segmentation. Even when domain-specific training provides limited improvement as a starting point for finetuning, the resulting model can segment unseen root datasets without any target annotations, reaching 99.7% of fine-tuned Dice on Sesame and exceeding fine-tuned on Switchgrass. In some cases, zero-shot segmentation provides root length measurements that match or even surpass the fine-tuned model in terms of root length correlation with ground truth annotations (Table VII). This has practical value when annotation data or training resources are unavailable: no labelled images are needed, no model training is required, and inference can run on an ordinary laptop with a simple GUI. For researchers with a new imaging setup, our root foundation model provides a working segmentation tool immediately.

### Corrective annotations may better reveal condition differences

DeepRootLab consistently shows the largest spread between conditions across both architectures (see Supplementary Figure 1). This may reflect its annotation approach: annotations were created by correcting model predictions rather than tracing roots independently. The annotations are concentrated on regions where the model makes errors, making them more direct indicators of reducible model weaknesses, and the corrective annotation protocol [Smith et al., 2022] encourages annotators to label only regions where they are confident, likely reducing label noise. Together, these properties likely allow the corrective annotations to serve as a more sensitive evaluation of model quality, making performance differences between training conditions more apparent on this dataset.

### Limitations

One possible explanation for the small full-data benefit of root foundation training is catastrophic forgetting: each fine-tuning stage may overwrite knowledge from the previous one rather than building on it [Banet et al., 2024, Ramasesh et al., 2021].

All conditions used full fine-tuning at every stage. Parameter-efficient fine-tuning may better preserve knowledge from earlier training stages, particularly in smaller models, and could yield a larger full-data benefit from root foundation training.

Root foundation training did not produce a statistically significant improvement on either architecture. Evaluating additional architectures pre-trained on different datasets would help clarify when domain-specific foundation training is worthwhile.

We used supervised training, limiting the foundation model to annotated datasets. Self-supervised pre-training would allow much larger collections of unannotated root images to be used, potentially providing stronger domain-specific representations.

For the few-shot experiments, a full validation and test set was used to facilitate comparison on the same images, but in practice having a smaller validation set corresponding to the limited training set may further degrade performance.

### Model release

We release our trained MobileSAM root foundation model for use with RootPainter (version 0.3.0 or later; https://github.com/Abe404/root_painter/releases) [Smith et al., 2022], enabling fully automatic root segmentation on new datasets with an ordinary laptop or desktop computer, with no need for annotation or training. The model weights are available on Zenodo at https://doi.org/10.5281/zenodo.20180274. Step-by-step instructions for installation and use are provided in the Supplementary Material. Segmentations produced by the released model can be passed through RhizoVision Explorer [Seethepalli et al., 2021] to extract per-image total root length and other traits, providing a fully automated GUI-based pipeline from raw images to root traits.

## Conclusion

We trained and evaluated the first domain-specific foundation model for root segmentation, using a leave-onedataset-out design across nine diverse root datasets with two architectures. The domain-specific model segments unseen root datasets zero-shot at 92% of fine-tuned Dice on average, exceeding 90% for 5 of 9 datasets. Fine-tuning on only 10 patches, it recovers 95% of its full-data Dice on average, versus 69% for a general pre-trained model. With full target-data fine-tuning, the two perform comparably, with mean improvements of +0.011 Dice for MobileSAM and +0.022 for M2F Swin-S, neither significant (Wilcoxon *p* = 0.150 and 0.064).

## Funding

The work presented in this article is supported by Novo Nordisk Foundation grant [NNF22OC0080177] for Abraham George Smith and Jens Petersen.

## Notes

https://doi.org/10.5281/zenodo.20180274

https://github.com/Abe404/root_painter/releases

https://github.com/sotlampr/seg

### Competing Interest Statement

The authors have declared no competing interest.

### Summary of Updates

This v2 revision is a presentation and language update. No reported numbers, methods, or conclusions are changed. Abstract: Restructured to lead with the 92 percent zero-shot retention result, then few-shot performance, then full fine-tuning. Added a sentence noting that the general pre-trained model often failed to converge at low patch counts (5 of 9 datasets gave Dice below 0.05 at 3 patches), while the root foundation model produced Dice above 0.47 on every dataset and patch count. Introduction: Added four citations to support the claim that each new imaging setup typically requires training a new model on an annotated subset of experimental data. Results: Within-section reordering so that prose now appears before the table that supports it. The Zero-shot performance subsection was moved earlier, to sit inside the H2 flow. The qualitative figure was moved earlier so it sits alongside the main results. Figures: Plots regenerated as vector PGF graphics for cleaner rendering. No changes to reported Dice values, deltas, p-values, methodology, dataset list, or release artifacts. The released MobileSAM root foundation model on Zenodo (10.5281/zenodo.20180274) and the public code repo at https://github.com/Abe404/RootFoundation are unchanged.

