## supplement for "A Root Foundation Model for Zero-Shot Segmentation"

### Supplementary Material: A Root Foundation Model for Zero-Shot Segmentation

#### Using the Foundation Model with RootPainter

The pre-trained MobileSAM root foundation model can be used to segment root images using the RootPainter software. No programming or command-line experience is required. The following instructions describe how to install RootPainter, download the foundation model, and segment a folder of images.

##### *Step 1: Install RootPainter*

We recommend the **Workstation edition**, which bundles both the GUI client and the trainer server in a single self-contained application. Download the installer for your operating system from the GitHub releases page:

https://github.com/Abe404/root_painter/releases

- **macOS:** Download and open the `.pkg` file. Follow the installer prompts. Since the application is not signed with an Apple Developer certificate, macOS Gatekeeper may block it. To proceed: right-click the installer and select **Open**, then go to **System Settings → Privacy & Security** and click **Open Anyway**.
- **Windows:** Download and run the `.exe` installer. Windows SmartScreen may show a warning for unrecognized applications. Click **More info** and then **Run anyway** to proceed.
- **Linux (Ubuntu):** Download the `.AppImage` file and make it executable (`chmod +x`).

Version 0.3.0 or later is required for MobileSAM support.

Alternatively, if you wish to use a remote GPU, the trainer and client can be installed separately. See the RootPainter documentation for details: https://github.com/Abe404/root_painter

##### *Step 2: Set Up the Sync Directory*

When you first open RootPainter, you will be asked to specify a *sync directory*. This is a folder on your computer where RootPainter stores its data. Choose or create an empty folder (e.g. `root_painter_sync` on your desktop).

##### *Step 3: Download the Foundation Model*

Download the pre-trained MobileSAM foundation model from Zenodo: https://doi.org/10.5281/zenodo.20180274. The file is named `mb_sam_vit_t_foundation_allin_pretrained_checkpoint_best.pth` and is 39 MB. Place this file somewhere inside your sync directory, for example in a `models` subfolder:

```
root_painter_sync/
    models/
        mb_sam_vit_t_foundation_allin_pretrained_checkpoint_best.pth
```

##### *Step 4: Segment Images*

1) Open RootPainter.
2) From the menu bar, select **Network → Segment folder**. In the workstation edition, this will automatically start the trainer server if it is not already running.
3) In the Segment Folder dialog:
   - Click **Specify input directory** and select the folder containing your root images.
   - Click **Specify output directory** and select (or create) a folder where the segmentation masks will be saved.
   - Click **Specify model file** and select the .pth file from Step 3.
   - Choose the desired output format (the default PNG with alpha channel is recommended for visual inspection).
4) Click **Segment**. A progress dialog will appear while the images are being processed.

The output segmentation masks will be saved in the specified output directory, one per input image. On a machine with a GPU (CUDA or Apple Silicon), segmentation takes approximately 1–2 seconds per image. CPU-only machines will be slower but fully supported.

##### *Notes*

- The foundation model performs dense root segmentation without any user annotation or fine-tuning. It can be applied directly to new root images.
- The model and dataset files must be located inside the sync directory for RootPainter to access them.
- The model was trained on 1024×1024 tiles; larger images are automatically tiled and reconstructed.
- RootPainter also supports interactive annotation and training of custom UNet models. See the main RootPainter documentation for details: https://github.com/Abe404/root_painter

#### Dice across conditions

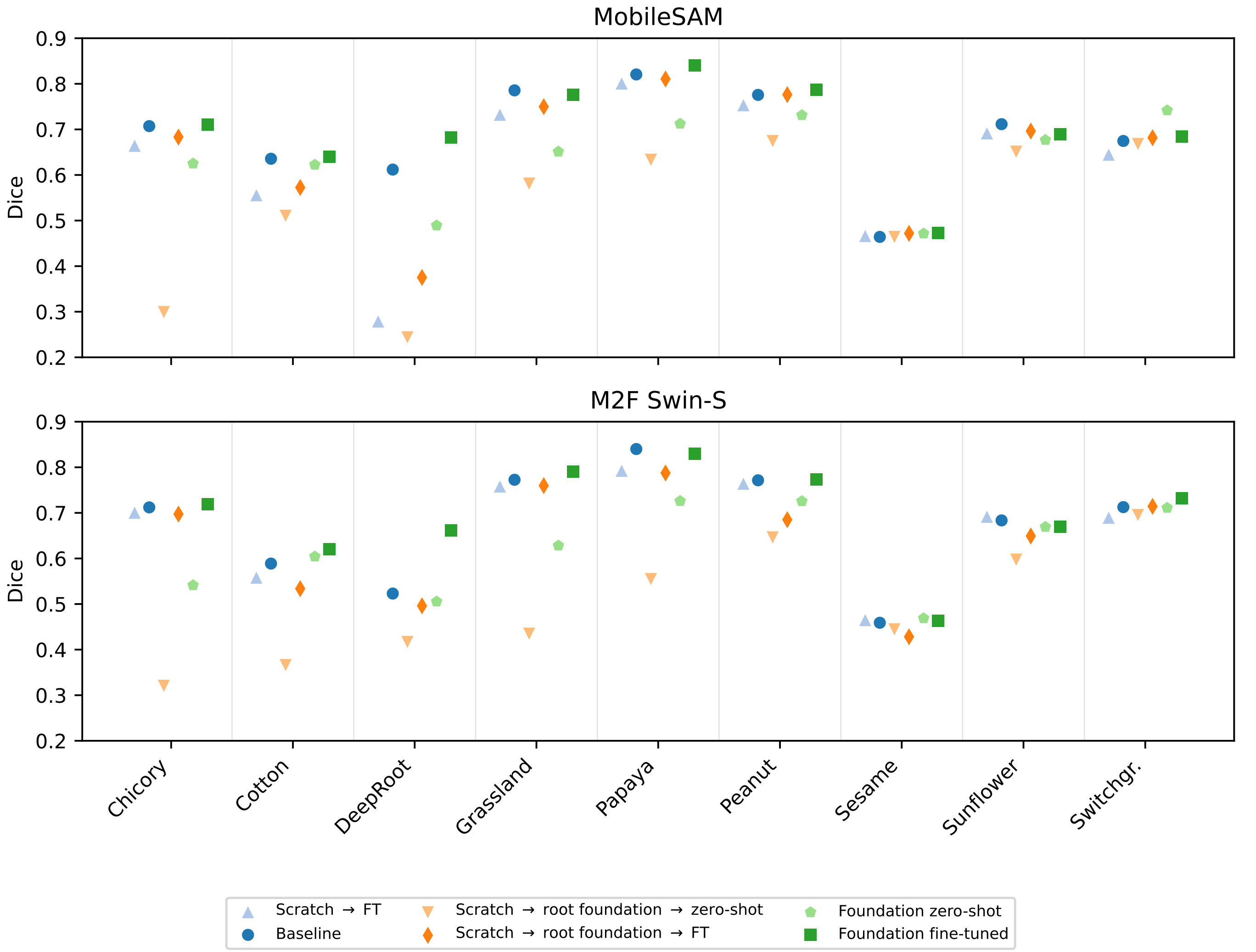


Fig. 1: Per-dataset Dice scores for all experimental conditions (test split). Each column is one held-out dataset; markers show the conditions from Table I.

#### Per-dataset results

TABLE I: Per-dataset Dice scores for MobileSAM (ViT-T), test split.

| Condition | Chicory | Cotton | DeepRoot | Grassland | Papaya | Peanut | Sesame | Sunflower | Switchgr. |
|---|---|---|---|---|---|---|---|---|---|
| pretrained zeroshot | 0.625 | 0.622 | 0.489 | 0.651 | 0.712 | 0.731 | 0.472 | 0.677 | 0.742 |
| pretrained finetuned | 0.711 | 0.640 | 0.682 | 0.776 | 0.840 | 0.787 | 0.473 | 0.690 | 0.684 |
| baseline pretrained | 0.707 | 0.636 | 0.612 | 0.786 | 0.821 | 0.776 | 0.464 | 0.711 | 0.675 |

TABLE II: Per-dataset Dice scores for M2F (Swin-S), test split.

| Condition | Chicory | Cotton | DeepRoot | Grassland | Papaya | Peanut | Sesame | Sunflower | Switchgr. |
|---|---|---|---|---|---|---|---|---|---|
| pretrained zeroshot | 0.541 | 0.604 | 0.506 | 0.628 | 0.726 | 0.726 | 0.469 | 0.669 | 0.711 |
| pretrained finetuned | 0.719 | 0.620 | 0.662 | 0.791 | 0.829 | 0.773 | 0.463 | 0.670 | 0.732 |
| baseline pretrained | 0.712 | 0.589 | 0.523 | 0.773 | 0.840 | 0.771 | 0.459 | 0.683 | 0.713 |
